# Anion supercharging enables structural assignment of oxidatively released *O*-glycans

**DOI:** 10.1101/2025.11.23.689986

**Authors:** Maia Kelly, Christopher Ashwood

## Abstract

*O*-glycan analysis faces analytical challenges due to poor fragmentation characteristics of oxidatively released *O*-glycan acids, which can preserve labile *O*-acetyl modifications but suffer from charge fixation at the reducing end. This study introduces a supercharging approach using hexafluoroisopropanol (HFIP) and butylamine mobile phase additives for enhanced mass spectrometric analysis of bleach-released *O*-glycan acids.

HFIP/butylamine increased average charge states by up to 50% compared to traditional mobile phases, dramatically improving MS2 fragmentation quality and enabling discrimination between isomeric compositions and structures. Enhanced fragmentation enabled confident identification of sialic acid variants and acetylation patterns, overcoming the analytical bottleneck of poor *O*-glycan acid fragmentation.

Computational optimization reduced database search times by up to ninefold while maintaining sensitivity. Application to multi-species gastric mucins revealed distinct glycosylation profiles. Porcine mucin showed predominant serine-linked neutral structures, while bovine and ovine mucins exhibited primarily threonine-linked sialylated glycans. A minor fraction of porcine and bovine mucins contained acetylated serine-linked sialylated glycans. This methodology provides a comprehensive framework for *O*-glycan characterization while preserving biological modifications.

## Introduction

Glycosylation, the covalent conjugation of sugars to biomolecules, protects glycoconjugates from degradation^1^ and provides ligands for sugar-specific binding^2^. For proteins, mucin domains are often heavily and densely modified with sugars, playing a critical role in maintaining the protective mucus barrier of the stomach^3^ and offering ligands for microbial communities^4^. The glycosylation status of gastric mucins determines which bacterial species can successfully adhere and establish residence,^5^ while simultaneously modulating immune recognition and the overall integrity of the gastric mucosal defense system^6^, making glycome-microbiome interactions crucial in both disease pathogenesis and therapeutic targeting^7^.

The detection and characterization of glycoconjugates in complex biological samples is fundamental for identifying the specific structural aberrations and linking them to pathological discrepancies with downstream implications for biomarker discovery. The release of glycans from their protein backbone decreases the complexity introduced by the protein component and allows for analytical methods that can be better tailored towards glycan-specific analysis. This approach is specifically focused on identification of structural differences^8^. The advantage of this approach is through the simplification of the analytical workflow and the further enhancement for structural resolution and sensitivity of glycan detection methods.

Current methodological approaches for *O*-glycan analysis face significant technical limitations owing to aiming for comprehensive release while retaining labile modifications. While the number of mucin proteases are increasing, such as StcE^9^and IMPa^10,11^, enzymes available for the comprehensive release of *O*-glycans are largely undescribed. The notable exception is specialized enzymes that can release de-sialylated *O*-glycans^12^ or recently, any form of *O*-glycan that modifies an accessible peptide sequence^13^. This enzymatic limitation contrasts sharply with *N*-glycan analysis, where well-characterized enzymes like PNGase F are routinely employed and are active on all mammalian *N*-glycans^14^. Following release, glycans are often subsequently reduced by sodium borohydride in an alkaline solution to produce peeling-resistant alditols that achieve higher chromatographic resolution^15^. However, alkaline solutions result in the loss of base-labile substituents such as *O*-acetyl esters, which can be biologically relevant^16^. Therefore, a mild *O*-glycan release method that does not result in peeling and is compatible with labile substituents is essential for comprehensive *O*-glycome analysis, particularly relating to *O*-acetylation.

One mild method for *O*-glycan release involves hypochlorite, which results in the selective formation of glycolic or lactic acid glycosides, for serine or threonine *O*-linked glycans, respectively^17,18^. This method retains the unique information about the modified amino acid that is typically lost during release, while simultaneously locking the glycan in a closed ring configuration.

This modification prevents destructive peeling reactions that can generate artifactual structures that are not originally present in the sample. This chemical preservation strategy maintains both structural integrity and linkage-specific information that is crucial for biological interpretation of labile *O-*glycan substituents.

The retention of *O*-glycan amino acid specificity through the formation of amino acid specific glycolic or lactic acid glycosides provides additional confidence in the identification of glycans as true *O*-glycans. In the case of *O*-GalNAc glycosylation, only serine and threonine residues can be modified in mammalian systems^19^, compared to free oligosaccharides or *N*-glycans which can have similar or identical glycan compositions but exist in different glycoconjugate forms. This specificity is particularly valuable in complex biological samples where multiple glycan types may co-exist. There is also the potential for enhanced analytical specificity (theoretically, a 2-fold increase in analyte specificity) as the *O*-glycan composition is modified with a mass indicative of the conjugated amino acid, simultaneously providing both glycan and amino acid information.

Mass spectrometric analysis of the modified glycan species presents additional analytical challenges due to their unique reducing end modifications. These modifications may alter charge distribution, modifying fragmentation patterns, however these fragmentation patterns can be partially reverted by supercharging^20^. Supercharging reagents are chemical additives used in electrospray ionization mass spectrometry (ESI-MS) to enhance the charge states of analyte ions, thereby improving sensitivity, resolution, and fragmentation efficiency for structural characterization^21^. These reagents function by altering the surface tension and pH of electrospray droplets, promoting more efficient charge transfer and increasing the average charge state of multiply charged ions. The enhanced charging not only improves signal intensity but also facilitates more extensive fragmentation in tandem MS experiments owing to fewer neutral (undetectable) fragments, leading to more confident structural elucidation^22^. For negative ion mode applications, commonly used supercharging reagents include meta-nitrobenzyl alcohol, benzyl alcohol, and various organic acids such as acetic acid and formic acid^23^.

We reveal that the produced *O*-glycan acid has remarkably poorer fragmentation compared to their unmodified *O-*glycan counterpart. This leads to significant impairment for compositional confirmation and structural elucidation capabilities. This fragmentation deficiency represents a critical analytical bottleneck that limits the comprehensive characterization of the produced *O*-glycan acid structures by LC-MS. Through in-depth analysis of three mobile phase additives, ammonium bicarbonate, ammonium fluoride, and a combination of hexafluorisopropanol (HFIP)/butylamine, for negative mode ionization, a mobile phase with HFIP and butylamine have been identified as an effective supercharging reagent pair. Along with modification of *O*-glycans with their corresponding acids, the utilization of supercharging drastically improves product ion diversity and therefore enhances structural elucidation capabilities, representing a significant methodological advancement for *O*-glycan analysis.

The improved analytical capabilities for structural elucidation combined with bioinformatic improvements to glycan database searching enable more confident and comprehensive glycan identification from complex mass spectrometric data. This work applies the optimized methodology to bovine, ovine, and porcine gastric mucins, systematically contrasting the *O*-glycan profiles across these three species to provide insights into species-specific glycosylation patterns, revealing differing ligands available for microbial interaction.

## Results and discussion

### O-glycan acid fragmentation patterns differ from released O-glycans

The oxidative release process modifies typical free *O*-glycans by retaining a sugar acid fragment derived from the amino acid component. This reducing-end modification alters analytical properties including monoisotopic mass and retention time, while simultaneously providing additional information about the specific amino acid that was modified with the respective *O*-glycan.

*O*-glycans released from serine residues retain a glycolic acid fragment, while those released from threonine retain a lactic acid fragment. Recent work in this area has underutilized anion MS2 analysis in their methodologies and without inclusion of MS2 analysis^17^, compositional confirmation and further structural elucidation capabilities can be hindered.

In **Figure 1**, oxidatively released *O*-glycan acids, retaining the respective sugar acid from the serine and threonine, exhibit distinct fragmentation patterns compared to their non-modified counterparts. These modified *O-*glycan acids yield limited structural detail for terminal fragments from the increase of the Y_n_ ions due to charge localization at the reducing end. This fragmentation limitation is particularly evident when attempting to distinguish *N*-glycolylneuraminic acid (NeuGc) from *N*-acetylneuraminic acid (NeuAc) (**Figure 1A**) and confirming sialic acid acetylation (**Figure 1B**). While diagnostic fragments containing sialic acid moieties are partially observed, they occur at levels decreased by 20-fold or greater than those observed for free *O*-glycans.

**Figure 1.**
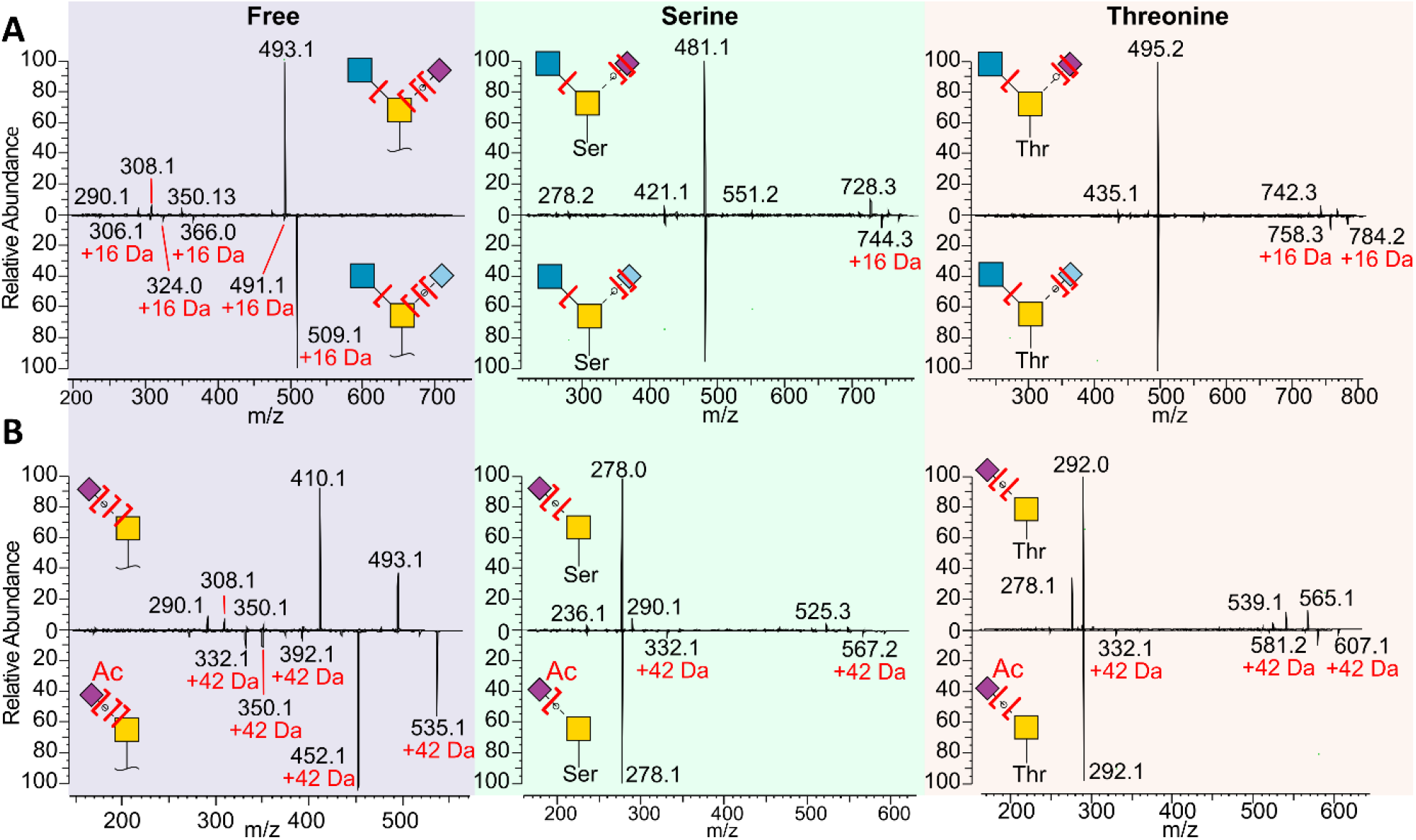
Oxidatively released *O*-glycan fragmentation spectra is less structurally informative compared to free *O*-glycans. Comparison of singly charged MS2 spectra for free, serine, and threonine-linked *O*-glycans for **A** NeuAc and NeuGc modified (16 Da difference). **B** Acetylated NeuAc (42 Da difference). Fragments retaining the compositional difference between the two structures are highlighted in red.

Addressing these MS2 analytical limitations is particularly crucial for acetylated *O*-glycan analysis, which represents a key analytical advantage and strength of the oxidative release methodology compared to alkaline conditions used in reductive amination and glycan reduction^14^.

### Screening of anion supercharging reagents for LC-MS

The oxidative release of the *O*-glycans lead to a shift from charge localization on the sialic acid to the reducing end sugar acid left from the serine or threonine acid counterpart. We hypothesized that increasing glycan charge state would overcome the charge fixation on the reducing end and lead to increased MS2 fragmentation and quality with corresponding improved structural coverage, similar to Huang et al when studying sulphated glycosylaminoglycans^24^. Three different LC-MS buffer compositions were evaluated to compare their effects on average charge state for each glycan species (**Figure 2A**). These included the traditional ammonium bicarbonate mobile phase additive, an LC-compatible anion supercharger ammonium fluoride^25,26^, and our recently applied HFIP/butylamine additive^27^.

**Figure 2.**
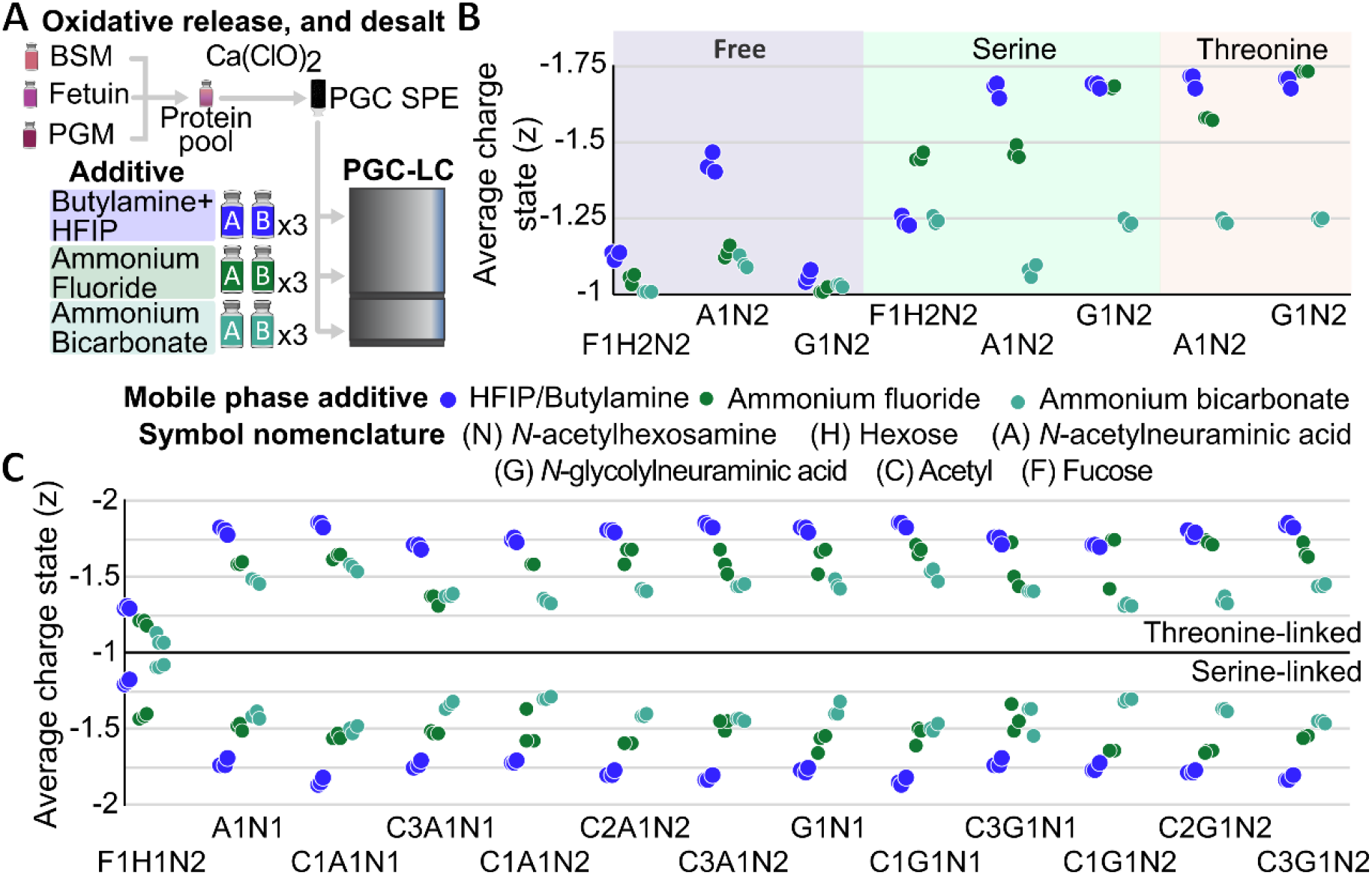
Mobile phase composition affects *O*-glycans charge state with HFIP/Butylamine identified as a supercharger. **A** Experimental setup for comparison of each mobile phase additive. Average observed charge state for shared glycans between **B** Free, Serine-linked, and Threonine-linked *O*-glycans or **C** Serine-linked and Threonine-linked *O*-glycans

For free *O*-glycans, notable average charge state increases were observed across the different mobile phases including a 35% increase in charge state for the A1N2 glycan composition (**Figure 2B**). When comparing the same mobile phases across free *O*-glycans, serine (glycolic acid) and threonine (lactic acid) acids, the *O*-glycan acids demonstrated greater susceptibility to charge state enhancement, with the same glycan composition (A1N2) exhibiting charge state increases exceeding 50%. Additionally, oxidative release at pH 7 enabled the detection of sialic acid *O*-acetylation, a labile modification lost at high pH^28–30^. Across all glycan compositions tested, with the exception of F1H1N2, the HFIP/butylamine buffer system consistently produced the highest charge states, confirming its effectiveness as a supercharger for negative mode mass spectrometry analysis (**Figure 2C**).

While we have shown the supercharging effects for *O*-glycan and *O*-glycan acids, we anticipate similar effects for other forms of glycosylation including free oligosaccharides, polysaccharides, and *N*-glycans. Furthermore, the moderate flow rates used here (150 µL/min) are higher than those traditionally used for supercharging studies, which rely on nanospray at nL/min flow rates, which may alter the supercharging observed here^31^.

### Accurate precursor m/z is insufficient for glycan composition verification

Software-aided analysis of the supercharged *O*-glycans led to putative identifications of many isomeric and isobaric *O*-glycan acids. However accurate assignment of acetylated glycan compositions between isomers and isobars was challenging due to the similarity in mass in combination with other monosaccharides. These compositional ambiguities frequently necessitated MS2 verification to ensure correct assignment. Specific isomeric challenges observed in this study are comprehensively detailed in **Table 1**, which includes chemical formulas, mass offsets, and our applied analytical solutions for each case.

**Table 1.** Isomeric and isobaric glycan motifs encountered during oxidative release of *O*-glycans.

| Glycan motif and mass 1 | Glycan motif and mass 2 | Chemical formula(s) | Mass offset (Da) | Identified solution |
| --- | --- | --- | --- | --- |
| (dHex)<sub>1</sub> (Ser)<sub>1</sub><br>204.063 | (Hex)<sub>1</sub> (Acetyl)<sub>1</sub><br>204.063 | C<sub>8</sub>H<sub>12</sub>O<sub>6</sub> | 0, isomeric | MS2 |
| (NeuAc)<sub>1</sub> (Ser)<sub>1</sub><br>349.101 | (NeuGc)<sub>1</sub> (Acetyl)<sub>1</sub><br>349.101 | C<sub>13</sub>H<sub>19</sub>NO<sub>10</sub> | 0, isomeric | MS2 |
| (Hex)<sub>1</sub> (HexNAc)<sub>1</sub> (Thr)<sub>1</sub><br>455.163 | (dHex)<sub>1</sub> (NeuAc)<sub>1</sub><br>455.163 | C<sub>17</sub>H<sub>29</sub>NO<sub>13</sub> | 0, isomeric | MS2 |
| (Hex)<sub>1</sub> (NeuAc)<sub>1</sub><br>453.148 | (dHex)<sub>1</sub> (NeuGc)<sub>1</sub><br>453.148 | C<sub>17</sub>H<sub>27</sub>NO<sub>13</sub> | 0, isomeric | MS2 |
| (NeuGc)<sub>1</sub> (Acetyl)<sub>1</sub><br>349.101 | (dHex)<sub>1</sub> (HexNAc)<sub>1</sub><br>349.137 | C<sub>13</sub>H<sub>19</sub>NO<sub>10</sub><br>C<sub>14</sub>H<sub>23</sub>NO<sub>9</sub> | 0.036 | High resolution MS1<br>MS2 |
| (Sulf)<sub>2</sub> (Ser)<sub>1</sub><br>217.919 | (dHex)<sub>1</sub> (Thr)<sub>1</sub><br>218.079 | C<sub>2</sub>H<sub>2</sub>O<sub>8</sub>S<sub>2</sub><br>C<sub>9</sub>H<sub>14</sub>O<sub>6</sub> | 0.160 | High resolution MS1<br>MS2 |

### Higher charge states improve isomeric structure discrimination by MS2

As no oxidative release conditions have been identified that completely prevent the co-generation of free *O*-glycans alongside the desired *O*-glycan acids, these findings demonstrate the critical importance of high-quality MS2 analysis for confirming that glycan compositions match expected structures (**Figure 3A**). While alternative orthogonal analytical methods such as ion mobility spectrometry or the development of comprehensive retention time libraries could potentially address these compositional challenges such as those by Vos *et al*^*32*^, our approach also enables characterisation of previously undescribed structures.

**Figure 3.**
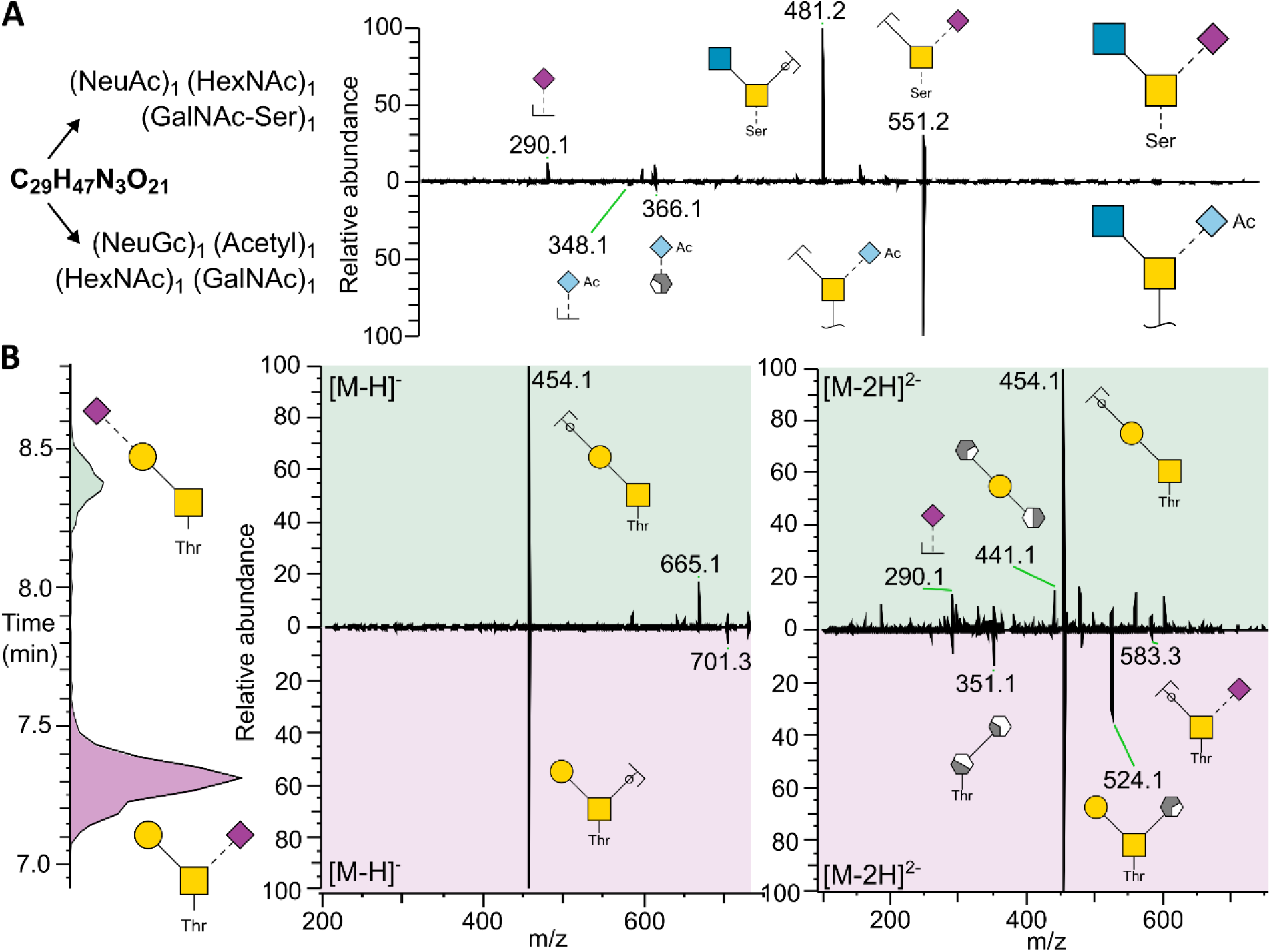
Supercharging overcomes reducing end charge fixation of *O*-glycan-acids enabling composition confirmation and isomer discrimination **A** Singly charged compositional isomer MS2 are largely identical **B** Higher charge states overcome poor MS2 spectrum quality

The necessity for high quality MS2, demonstrated by isomeric glycan compositions (**Table 1**), is reinforced by singly charged MS2 spectra that are essentially identical across isomeric structures. The only distinguishing feature in these singly charged spectra may be the minor presence of potentially discriminative product ions, such as the ion observed at *m/z* 665.1 (**Figure 3B**), which provides insufficient structural information for confident isomer identification. In contrast, higher charge states, exemplified by doubly charged precursor ions, generate distinctly different fragmentation patterns between isomeric structures. These enhanced charge states enable clear characterization of glycosidic bonds and linkage positions, providing structural detail for isomer identification and deep glycan characterization.

In addition to the higher charge state and more diverse fragmentation patterns, resonant CID also benefits from the lower precursor *m/z*, enabling retention of product ions often lost to the 1/3^rd^ rule for ion trap mass spectrometers^33,34^, further improving confidence in compositional annotation. While spectral library-based approaches, such as Unicarb-DB^35,36^, can be leveraged for glycan discovery, these vastly different MS2 spectra between *O*-glycans and *O*-glycan acids necessitates extension of these libraries (**Figures 1 and 3**). Future research in this area, based on the results shown here, may enable these libraries to be extended in-silico.

### Computational optimisation improves data analysis throughput

With the analytical methodology established for bleach-released *O*-glycan acids, one final challenge remained: the rapid assignment of glycan compositions to MS2 spectra while prioritizing composition verification and structural assignment. This step is crucial for efficient workflow implementation in high-throughput glycan analysis. As *O*-glycans can contain multiple unique monosaccharide building blocks, their comprehensive inclusion in database searching is essential for achieving high analytical coverage. Each additional monosaccharide incorporated into the search space exponentially increase computational search times when using GlyCombo, a glycan composition search engine (**Figure 4A**). This computational burden presented a significant bottleneck for practical application of the methodology, particularly when encountered *O*-glycans that can have three different reducing ends: free, serine-based (glycolic acid), and threonine-based (lactic acid) *O*-glycan acids.

**Figure 4.**
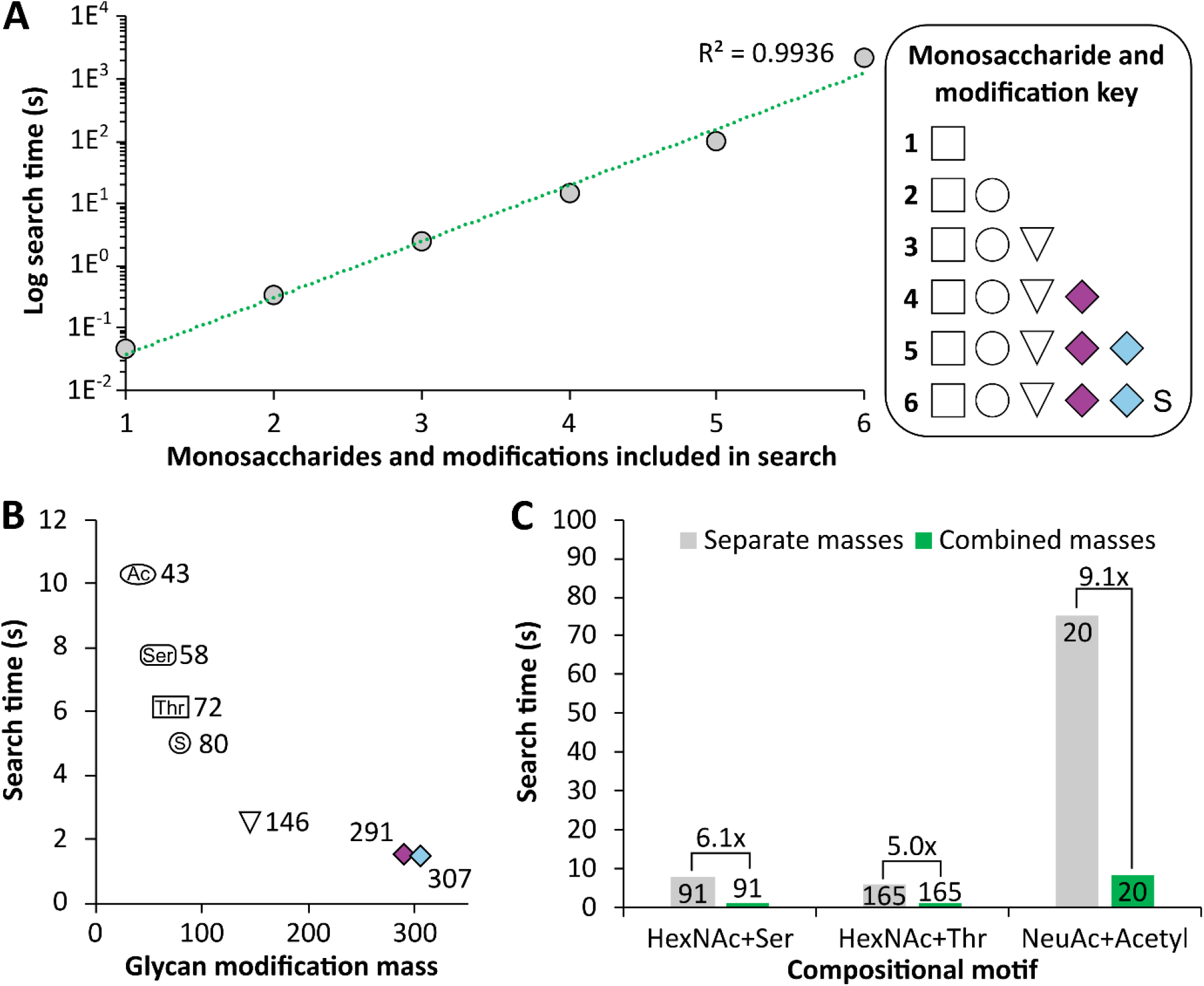
Searching modified monosaccharides rapidly assigns bleach released supercharged *O*-glycans to PGM *O*-glycan data. **A** Search time increases exponentially with monosaccharide and modification count. **B** Search time is inversely proportional to modification mass. **C** Combining compositional motifs result in drastically reduced search times with no reduction in number of matches. Column labels correspond to the number of composition matches for each approach.

For the analytical approach used in this study, where base-labile *O*-acetyl groups are preserved and *O*-glycan acids are specifically required for database searching, the increased search time associated with their inclusion showed an inverse correlation with molecular mass. Specifically, modifications with higher molecular masses resulted in proportionally smaller increases in search time (**Figure 4B**), suggesting that computational efficiency could be optimized by strategic grouping of modifications as a custom modification based on known pairings.

Many modifications occur on specific monosaccharide residues and the low mass nature of many modifications exacerbates the computational search time. Combining low mass modifications such as acetylation, serine *O*-glycan acids, and threonine *O*-glycan acids, with their modified monosaccharide residue allowed us to develop a strategy to drastically reduce the computational impact of low mass modifications. This optimization strategy achieved up to a nine-fold improvement in search time while maintaining the same number of compositional matches containing the target compositional motifs, demonstrating no loss in sensitivity (**Figure 4C**).

### Application of supercharging to gastric mucin across three mammalian species

This computational enhancement enabled the practical application of the methodology to large-scale datasets, including comprehensive multi-species gastric mucin comparative studies that would have been computationally prohibitive using conventional search strategies. Structural elucidation was successfully achieved through the implementation of supercharged PGC-LC-MS analysis, with effective isomeric separation demonstrated in **Figure 5A**. This example illustrates that while porcine gastric mucin (PGM) contained five distinct isomeric structures, only one representative structure from each isomeric group was detected with high confidence in ovine gastric mucin (OGM) and bovine gastric mucin (BGM). This analytical approach was subsequently extended to analyze all *O*-glycan acids observed across the three sample types, with quantitative assessment of each identified structure.

**Figure 5.**
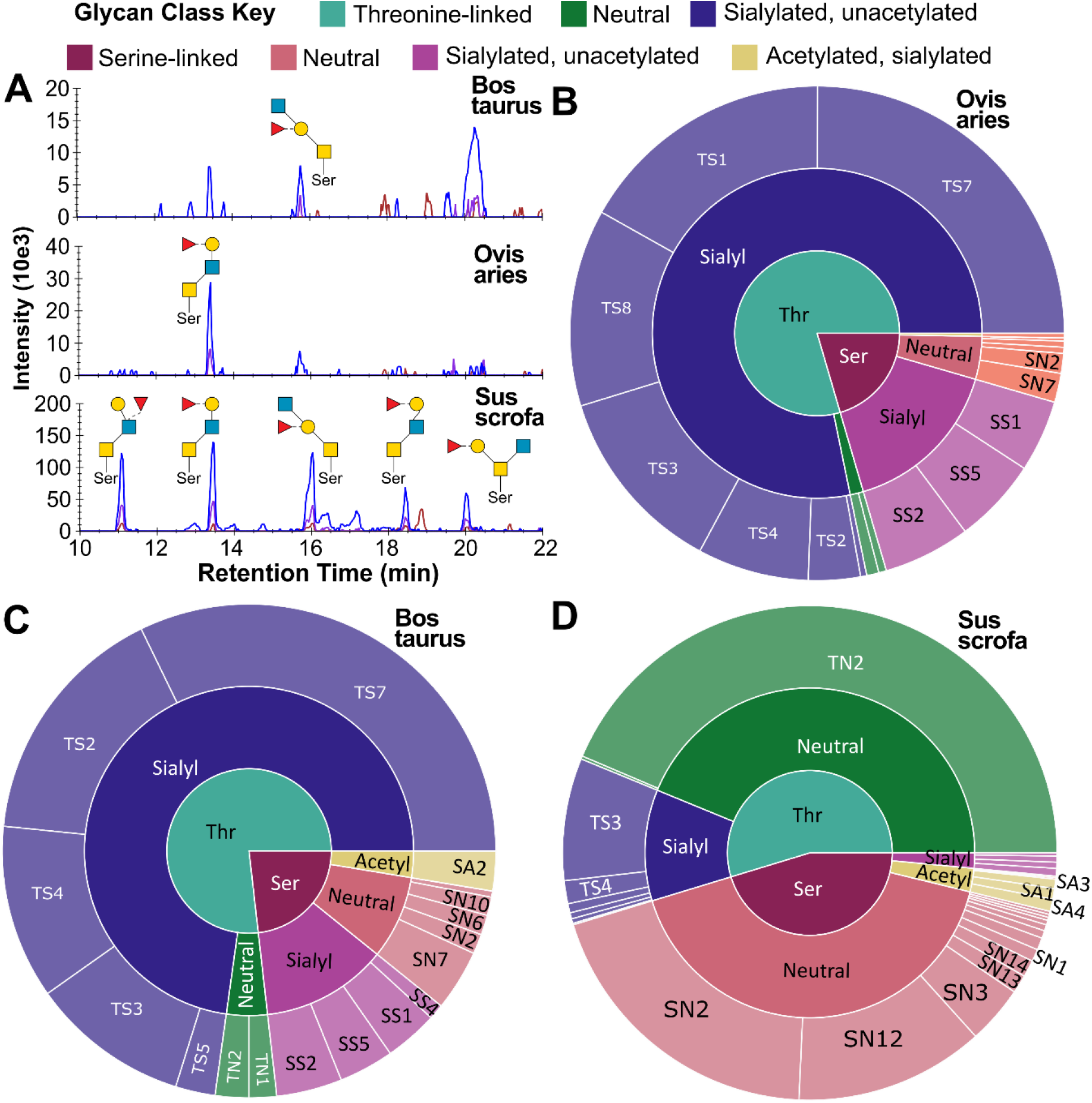
Supercharged *O*-glycomics enables multi-species comparisons of gastric mucin, revealing species-specific compositions and structures. **A** An example of isomeric diversity across the three species for a single glycan composition. **B-D** Sunburst charts of the *O*-glycan acids observed with respective linkage to serine or threonine for **B** ovine **C** bovine **D** porcine gastric mucins.

Ovine gastric mucin exhibited a glycan profile dominated by threonine-linked sialylated *O*-glycans, with the structural diversity largely distributed across six major glycan structures (**Figure 5B**). Notably, minimal acetylation was observed in the ovine samples. Bovine gastric mucin demonstrated a similar overall glycan profile to ovine mucin; however, a minor proportion of the glycan structures were acetylated, specifically featuring the SA2 structural variant, which differed from the acetylated structures observed in porcine samples (SA1, SA3, SA4) (**Figure 5C**).

Porcine gastric mucin displayed the most distinctive glycan profile among the three species examined. The most abundant glycan type was threonine-linked, neutral glycans (TN2), comprising more than 30% of the total observed profile and represented by a single predominant structure. Approximately 45% of the porcine glycan profile consisted of serine-linked structures, with acetylation modifications predominantly associated with serine-linked *O*-glycans rather than threonine-linked variants (**Figure 5D**).

The successful implementation of this enhanced computational approach opens new avenues for understanding mucin glycobiology across broader taxonomic groups. Future investigations could leverage this methodology to explore temporal changes in mucin glycosylation during disease progression, developmental stages, or environmental adaptations. Additionally, the scalability of this analytical framework positions it well for integration with emerging multi-omics approaches, potentially enabling comprehensive mapping of glycan-protein-microbiome interactions in complex biological systems. The species-specific patterns identified here warrant further functional validation studies to elucidate the biological significance of these structural differences in gastric protection and host-pathogen interactions.

## Conclusions

The application of HFIP/butylamine supercharging to oxidatively released *O*-glycan acids represent significant progress in analytical glycomics. While supercharging provided minor improvements for free *O*-glycans, the dramatic enhancement observed for *O*-glycan acids overcame critical analytical limitations and enabled deep structural elucidation. The structural analysis across bovine, ovine, and porcine mucins provides foundational data for developing retention time and fragmentation libraries to further increase the throughput and accessibility of future *O*-glycan acid characterization studies. The observed elution patterns, where threonine-linked structures consistently elute before their serine counterparts, establish predictable chromatographic behaviour that will enhance confidence in structural assignments. These analytical capabilities, combined with optimized compositional matching strategies that achieve up to nine-fold improvements in computational efficiency, establish a robust framework for high-throughput *O*-glycan analysis.

The comparative glycomic analysis of gastric mucins across these three species reveals distinct structural patterns that have implications for understanding host-microbiome interactions in gastric health and disease. The species-specific differences observed—particularly the abundant neutral threonine-linked glycans and extensive serine-linked acetylation in porcine mucins versus the sialylation-dominant profiles in bovine and ovine samples—likely influence bacterial adhesion patterns and colonization dynamics within the gastric environment^37^. These structural variations may explain species-specific susceptibilities to gastric pathogens such as Helicobacter pylori^38^, as bacterial adhesins exhibit selectivity for particular glycan motifs. The preservation of acetylation patterns through oxidative release methodology is particularly relevant, as these modifications are known to influence bacterial recognition and immune system activation, potentially serving as key determinants in the establishment of protective versus pathogenic microbial communities in the gastric mucosa.

## Methods

### O-glycoprotein sources

All chemicals and reagents were purchased from Millipore Sigma (Castle Hill, Australia) unless specified otherwise. Purified glycoproteins (bovine fetuin, bovine submaxillary mucin, porcine gastric mucin) were purchased from Millipore Sigma (Castle Hill, Australia).

Gastric mucins were extracted from sheep and bovine stomach lining, obtained from a local butcher (Wollongong, Australia). Extraction was performed as described by Nordman *et al*^*39*^ with modifications, frozen stomach lining (1 g) was thawed by the addition of ice-cold extraction buffer (6 M guanidinium chloride, 5 mM EDTA, and 10 mM sodium phosphate buffer, pH 6.5). The mucosa were dispersed with Dounce homogenizers and chilled overnight at 4 °C. Insoluble material was removed by centrifugation (16,000 g for 5 min) and re-extracted twice with extraction buffer.

### Oxidative glycan release

The purified glycoproteins and stomach lining were solubilised in 5% SDS and 100 mM TEAB. Proteins were reduced with 5 mM dithiothreitol for 30 min at 55 °C, alkylated with 10 mM iodoacetamide in the dark for 30 min at room temperature, and quenched with an additional 5 mM dithiothreitol for 15 min at room temperature. Glycans were released from glycoproteins and purified for LC-MS analysis as described in Ashwood *et al*^*40*^, with minor modifications. In brief, proteins were precipitated with methanol and phosphoric acid, and purified via DNA miniprep silica columns (Bioneer, Republic of Korea). Glycans were oxidatively released as described in Vos *et al*^*17*^, with modifications. Immobilised protein on the miniprep silica column were resuspended in 100 µL of 25 mg/ml of Ca(OCl)_2_^41^, adjusted to pH 7 with formic acid, and left to react at room temperature for 30 minutes.

Following glycan release, glycans were eluted from miniprep silica columns with 400 μL of 0.1% formic acid and then desalted using Supelclean ENVI-Carb SPE (100 mg). The desalting procedure involved conditioning with 400 μL acetonitrile/0.1% formic acid, equilibration with 1.5 mL water/0.1% formic acid, sample loading, desalting with 1.5 mL water/0.1% formic acid, and final elution with 400 μL 50:50 acetonitrile:water containing 0.1% formic acid. The desalted glycan solutions were then dried by centrifugal evaporation. Glycans were resuspended in ultra-pure water and then transferred into a 96 well PCR plate for injection.

### LC-MS setup

Glycans were separated with a Thermo Fisher Scientific Vanquish Horizon HPLC (San Jose, USA) and ionised into an Orbitrap IQ-X Tribrid mass spectrometer (San Jose, USA). A Thermo Fisher Hypercarb PGC column (Lithuania, 100 mm length by 1 mm internal diameter, 3 micron pore size), held at 90 °C, was used for all separations. Mobile phase A composed of water and mobile phase B composed of acetone with 5 mM HFIP and 5 mM butylamine added. During supercharging optimisation, 10 mM of ammonium bicarbonate or ammonium fluoride were added instead of HFIP/butylamine. LC separation was performed at 150 μL/min. Glycans were separated over a 30 min run, with 0-15% B over 23 min, 100% B held for 3 min, then 100% A for 4 min.

A Thermo Fisher Scientific Tribrid IQ-X was set to negative mode in DDA MS2 mode with a total cycle time of 1.3 seconds. ESI voltage was 3 kV, with sheath and auxiliary gases at 30 and 20 arbitrary units, respectively. Precursor spectra were collected across two acquisition regimes. The first, a full scan of 337 – 1700 *m/z* was collected in an Orbitrap at 60,000 resolution to hit an AGC target of 1.4e^6^ with a maximum inject time of 400 ms. The second, a multiplexed set of windows inspired by MAP-MS^42^, across 135-165 m/z, 169-211 m/z, and 215-233 m/z were collected in an Orbitrap and scanned together at 7,500 resolution to hit an AGC target of 5e^5^ with a maximum injection time of 11 ms.

Resonant CID fragment spectra were collected with an isolation width of 1.5 m/z, maximum injection time of 200 ms, and an AGC target of 1e^5^. The normalised collision energy was set to 38 and scanned at 0.6 m/z resolution in a linear ion trap. Dynamic exclusion was utilised, excluding each precursor 6 seconds after fragmentation. Charge state filtering was performed only for the multiplexed MS1 scan, specifically fragmenting charge states -2 to -4.

### Data analysis

LC-MS raw files were analysed by GlyCombo^43^ (v1.2) to assign glycan compositions to precursor m/z values, identify the most intense MS2 scans for each structure for annotation in GlycoWorkBench^44^, and construct the Skyline assay detailing *O*-glycan and *O*-glycan acid structures. mzML input was used with an error tolerance of 25 ppm, reducing end specified as free, derivatisation as native and adducts set as M-H^-^. Monosaccharide search space was: Hex 0-4, HexNAc 0-2, dHex 0-1, NeuAc 0-2, NeuGc 0-2, and three custom monosaccharides (GalNAc-Ser) and (GalNAc-Thr) at 0-1 of each, and (NeuAc-Acetyl) at 0-3. The monoisotopic mass and chemical formula were assigned for each modification as follows: GalNAc-Ser (C_10_H_15_N_1_O_7_, 261.08485 Da), GalNAc-Thr (C_11_H_17_N_1_O_7_, 275.10050 Da), and NeuAc-Acetyl (C_13_H_19_N_1_O_9_, 333.10598 Da). Additional acetylation was added post-search in Skyline and subsequently quality filtered. Search time comparisons were performed on a consumer-grade Lenovo laptop equipped with an AMD Ryzen 6 Pro 5850U CPU, 16 GB of RAM and a 500 GB SSD.

Skyline-daily^45,46^ (v25.1) integrated the first three isotopic peaks with mass analyser set to centroid at 15 ppm mass accuracy. These isotopic integrations were used to quality filter identifications (>=0.9 idotp) and quantify glycans. The idotp value of 0.9 was empirically selected to remove poor quality MS1-matches (caused by monoisotopic peak misassignment, incorrect charge assignment, and poor signal to noise ratios) while preserving high-quality matches^47^.

### Data availability

The raw MS glycomics data generated in this study have been deposited in GlycoPOST^48^ under accession code https://glycopost.glycosmos.org/entry/GPST000629. All raw data, spectral libraries, and processed Skyline documents are available on Panorama^49^(https://panoramaweb.org/SuperchargedGlycomics.url).

## Notes

### Competing Interest Statement

C.A. is the director of Protea Glycosciences, a company which provides fee-for-service glycomics assays, analytical standards, and software.

